# Heterotypic Influenza Infections Mitigate Susceptibility to Secondary Bacterial Infection

**DOI:** 10.1101/2022.04.12.488032

**Authors:** Ellyse M. Cipolla, Molin Yue, Kara L. Nickolich, Brydie R. Huckestein, Danielle E. Antos, Wei Chen, John F. Alcorn

**Affiliations:** Department of Pediatrics, UPMC Children’s Hospital of Pittsburgh, Pittsburgh, PA; Department of Immunology, University of Pittsburgh, Pittsburgh, PA; Department of Biostatistics, School of Public Health, University of Pittsburgh, Pittsburgh, PA

**Author notes:** Corresponding Author John F. Alcorn, PhD, 9127 Rangos Research Building 4401 Penn Ave., Pittsburgh, PA 15224. Author Contribution – EMC, WC, JFA conceived the study EMC, JFA obtained funding EMC, KLN, BRH, DA performed the experiments EMC, MY, JFA analyzed the data EMC, JFA drafted and edited the manuscript.

**Keywords:** – Lung, Pneumonia, Virus, Staphylococcus aureus, Super-infection

## Abstract

Influenza associated bacterial super-infections have devastating impacts on the lung and can result in increased risk of mortality. New strains of influenza circulate throughout the population yearly promoting the establishment of immune memory. Nearly all individuals have some degree of influenza memory prior to adulthood. Due to this we sought to understand the role of immune memory during bacterial super-infections. An influenza heterotypic immunity model was established using influenza A/PR/8/34 and A/X31. We report here that influenza experienced mice are more resistant to secondary bacterial infection with methicillin-resistant *Staphylococcus aureus* as determined by wasting, bacterial burden, pulmonary inflammation, and lung leak, despite significant ongoing lung remodeling. Multidimensional flow cytometry and lung transcriptomics revealed significant alterations in the lung environment in influenza-experienced mice compared with naïve animals. These include changes in the lung monocyte and T cell compartments, characterized by increased expansion of influenza tetramer specific CD8^+^ T cells. The protection that was seen in memory experienced mouse model is associated with the reduction in inflammatory mechanisms making the lung less susceptible to damage and subsequent bacterial colonization. These findings provide insight into how influenza heterotypic immunity re-shapes the lung environment and the immune response to a re-challenge event, which is highly relevant to the context of human infection.

## Introduction

Seasonal influenza infections annually account for significant amounts of hospitalizations as well as increased morbidity and mortality especially in the young, immunocompromised, and the elderly. The WHO estimates that yearly influenza infections led to 290,000-600,000 deaths globally prior to the COVID-19 pandemic (1). Influenza infections begin by primarily targeting epithelial cells as sites for viral replication (2). Once epithelial cells become infected by the virus a cascade of innate and adaptive immunological mechanisms is initiated resulting in clearance of the virus. This process of viral clearance creates a highly inflamed lung environment that can lead to excessive damage to the lung epithelium and an increased risk of development of a secondary bacterial infection and subsequent pneumonia (3–7). Influenza associated secondary bacterial infections heighten the risk of mortality as is evidenced by historical histology records from the 1918 influenza pandemic indicating the presence of bacterial infections in the lungs of approximately 95% of those who succumbed to infection (8, 9). Mouse models have provided evidence of dysregulated immune responses to the bacteria, which are thought to be caused by the preceding anti-viral response (10–20). These mechanisms present in the lung environment allow for influenza and bacteria to act in a synergistic way, with influenza promoting subsequent bacterial colonization and outgrowth (21). Due to the extensive damage that influenza associated secondary bacterial infections can have on the host lung it is imperative to understand the mechanisms that shape the host-pathogen response to these types of infections.

Influenza evades host immunity by readily mutating its surface proteins, neuraminidase (NA) and hemagglutinin (HA), in response to evolutionary pressure that in part is driven by host antibodies directed against HA and NA (22, 23). This results in the yearly emergence of novel influenza strains. Although the antibody response is at risk of being evaded by heterotypic influenza strains, the host also establishes a memory immune cell compartment distinct from the antibody response that is able to mobilize and respond against multiple influenza strains (22, 23). Both the innate and adaptive immune systems can produce memory immune cells against a variety of pathogens (24, 25). The importance of the immune response following priming and repeated exposure to antigens has been studied extensively at both the basic and translational level for vaccine and therapeutic purposes (26–29). There is evidence that priming by influenza reprograms immune cells over time, specifically myeloid cells to display enhanced antibacterial functions (30). Both CD8^+^ and CD4^+^ memory T cells have been shown to be of particular importance in the memory response to heterologous strains of influenza because they are able to react against internally conserved pieces of the virus that are less likely to undergo mutations driven by evolutionary pressure, ensuring quick mobilization and responses against infection (31–33). There are several memory T cell subsets: central memory (Tcm), effector memory (Tem), and resident memory (Trm). The different subsets of memory T cells are defined by their localization as well as surface marker expression. Due to their location within the lung at the site of infection, Trm cells have been implicated in the augmented clearance of viral respiratory infections including influenza (34). Nearly all humans have some level of pre-existing memory against influenza from early life exposures. Although memory cells are imperative for clearance of heterotypic viral strains, a fine balance in the inflammatory response must be achieved to ensure the least amount of damage to the lung tissue.

Currently, most of the studies in the field on influenza associated bacterial infections focus on the acute stage in naïve mice, whereas our study takes into account pre-existing influenza memory akin to human influenza immunity. To better understand the role of memory in influenza-associated bacterial infections, we used a mouse model to mimic a heterotypic influenza infection using H3N2 and re-challenging approximately two months later with H1N1 followed in six days by infection with methicillin-resistant *Staphylococcus aureus* (MRSA). Using spectral flow cytometry and transcriptomics we studied the role that antigen experience has on susceptibility to secondary bacterial infections. The model we used here is intended to provide a more clinically relevant representation of how heterotypic influenza strains infect the human population annually and how the immune response is shaped over time. These data provide insight into how viral infections reprogram the lung to respond to subsequent infections and the repercussions this has on susceptibility to secondary infections, further highlighting the importance of experienced cells in the response to lung pathogens.

## Materials and Methods

### Mouse Model and Sample Collection

On day zero, six to eight-week old male WT C57BL/6 mice (Taconic Farms, Germantown, NY) were infected with 10^5^ pfu of mouse adapted influenza A/X31 H3N2 or PBS vehicle. After 53-54 days the mice were re-challenged with 100 pfu of a heterotypic strain of mouse adapted influenza A/PR/8/34 H1N1. 6 days after influenza re-challenge (day 59/60) the mice were challenged with 5×10^7^ colony forming units (cfu) of USA300 MRSA suspended in PBS and harvested a day later. All infections were given via oropharyngeal aspiration. Mice were maintained under pathogen-free conditions at UPMC Children’s Hospital of Pittsburgh and all animal studies were conducted with approval from the University of Pittsburgh Institutional Animal Care and Use Committee. All studies used age- and sex-matched mice. Mice were euthanized via pentobarbital injection followed by exsanguination by severing the renal artery. No mice died prior to euthanasia.

### Bronchoalveolar Lavage Fluid Collection and Differential Cell Counting

Upon harvest, mice were cannulated and lavaged with 1ml of PBS for bronchoalveolar lavage fluid (BALF) collection. BALF was spun down and pelleted and supernatant was collected and stored for downstream analysis. Pelleted cells were treated with ACK lysing buffer (Gibco Fisher Scientific, Hampton, NH) to remove red blood cells. The cell pellet was then resuspended in 500μl of PBS and total cell count was determined by hemocytometer. 200μLs of resuspended cells were then concentrated on a microscope slide using a cytocentrifuge (ThermoFisher Scientific, Waltham, MA) and stained with Diff-Quik staining solution (Fisher Scientific, Hampton, NH) to determine monocyte, neutrophil, eosinophil, and lymphocyte counts.

### Bacterial Plating

Right upper lung lobes from mice were collected and homogenized in 1ml of PBS. After homogenization, 10-fold dilutions were dot plated on culture plates. Plates were then incubated at 37° Celsius overnight, and cfu was assessed by bacterial colony counting.

### Flow Cytometry

Mouse lungs were aseptically dissected using sterile scissors. Lungs were then digested for an hour at 37° Celsius in 1mg/ml collagenase media (DMEM Gibco Fisher Scientific, Hampton, NH). After an hour, lungs were mashed through 70 micron filters to obtain a single cell suspension. The single cell suspension was treated with ACK lysing buffer (Gibco Fisher Scientific, Hampton, NH) to remove red blood cells. After red blood cell lysis, cells were resuspended in PBS. Single cell suspensions were stained as follows for spectral flow cytometry analysis. For the T cell memory panel, cells were stained with anti-Cd45 (30-F11,BD Pharmingen, San Diego, CA), Cd4 (RM4-5,BD Biosciences, Franklin Lakes, NJ), Cd103 (M290,Invitrogen-ThermoFisher Scientific, Waltham, MA), Cd49a (HA31/8,BD Biosciences, Franklin Lakes, NJ), Cd69 (H1.2F3, BioLegend, San Diego, CA), Pd-l2 (TY25,BioLegend, San Diego, CA), Pd-l1 (10F.9G2,BioLegend, San Diego, CA), Cx3cr1 (SA011F11,BioLegend, San Diego, CA), Lag3 (C9B7W, BioLegend, San Diego, CA), Cd8 (53-6.7,Invitrogen-ThermoFisher Scientific, Waltham, MA), Cd11a (M17/4,BioLegend, San Diego, CA), Cd127 (SB/199,Invitrogen-ThermoFisher Scientific, Waltham, MA), Klrg1 (2F1,BD Biosciences, Franklin Lakes, NJ), Foxp3 (FJK-16s, Invitrogen-ThermoFisher Scientific, Waltham, MA), Tetramer (I-A(b) Influenza A NP 311-325 QVYSLIRPNENPAHK), Tim3 (RMT3-23,BioLegend, San Diego, CA), CD244.2 (m2B4(B6)458.1,BioLegend, San Diego, CA), Pd-1 (29F.1A12,BioLegend, San Diego, CA), Cd90.2 (30-H12,BD Biosciences, Franklin Lakes, NJ), Cd62L (MEL-14,BD Biosciences, Franklin Lakes, NJ), and Cd44 (IM7,BD Biosciences, Franklin Lakes, NJ). NP366 tetramers were obtained from the NIH tetramer core facility (Bethesda, MD) and were stained at 37° Celsius for 30 minutes prior to viability stain. For the myeloid panel, cells were stained with anti-CD45 (30-F11,Invitrogen-ThermoFisher Scientific, Waltham, MA), Pd-l2 (TY25,BioLegend, San Diego, CA), F4/80 (T45-2342,BD Biosciences, Franklin Lakes, NJ), Cd64a/b (X54-5/7.1,BD Biosciences, Franklin Lakes, NJ), Pd-l1 (10F.9G2,BioLegend, San Diego, CA), Cd103 (2E7,Invitrogen-ThermoFisher Scientific, Waltham, MA), Ly6c (HK1.4,BioLegend, San Diego, CA), Cd11b (M1/70,BD Biosciences, Franklin Lakes, NJ), MHC-II (M5/114.15.2,BD Biosciences, Franklin Lakes, NJ), Cd80 (16-10A1,BD Biosciences, Franklin Lakes, NJ), Cd86 (GL1,BD Biosciences, Franklin Lakes, NJ), B220 (RA3-6B2,Invitrogen-ThermoFisher Scientific, Waltham, MA), Tcrb (H57-597,BioLegend, San Diego, CA), SiglecF (1RNM44N, Invitrogen-ThermoFisher Scientific, Waltham, MA), Cd24 (M1/69,BioLegend, San Diego, CA), Pd-1 (RMP1-30,BioLegend, San Diego, CA), Cd244.2 (m2B4(B6)458.1,BioLegend, San Diego, CA), Nk1.1 (PK136,BioLegend, San Diego, CA), Cd11c (N418,Invitrogen-ThermoFisher Scientific. Waltham, MA), Tim3 (B8.2C12,BioLegend, San Diego, CA), and Arg-1 (A1exF5,Invitrogen-ThermoFisher Scientific, Waltham, MA). The viability dye, Zombie NIR (BioLegend, San Diego, CA), was used to exclude live cells from dead cells in both panels. Our mastermix for cell staining contained Super Bright Complete Staining Buffer (ThermoFisher Scientific, Waltham, MA) as well as True-Stain Monocyte Blocker (BioLegend, San Diego, CA) for the myeloid panel. Intracellular staining was performed at room temperature using the eBioscience™ Foxp3/Transcription Factor Staining Buffer Set (ThermoFisher Scientific, Waltham, MA) as directed by the manufacturer. All samples were run on the Cytek Aurora (Cytek Biosciences, Fremont, CA). Flow cytometric analysis was performed using FlowJo with UMAP (65) and FlowSOM (66) integrated plug-ins (Tree Star, Ashland, OR). Absolute cell counts were determined following manufacturer’s instructions using UltraComp eBeads^TM^ Plus Compensation Beads (Invitrogen-ThermoFisher Scientific, Waltham, MA). Gating strategies are outlined is Supplemental Figures 1 and 2.

### Histology

Left lung lobes from mice were inflated with and preserved in 10% neutral buffered formalin solution. Formalin fixed tissues were then transferred to 70% ethanol and shipped to StageBio (Mount Jackson, VA), where they were paraffin embedded and sectioned for histopathological analysis. Upon sectioning, hematoxylin and eosin (H&E) staining was performed. Histological scoring was performed on H&E stained slides using a scale from 1 to 4 with 1 being no damage and 4 being severely damaged. Cellular infiltration and tissue damage was assessed for the lung parenchyma, peribronchial, and perivascular regions. Scoring was performed sample blinded by two separate investigators.

### RNA extraction and qPCR

Mouse lungs were isolated and snap-frozen in liquid nitrogen or suspended in Allprotect Tissue Reagent (Qiagen, Hilden, Germany). RNA was extracted as directed using the Qiagen RNeasy Mini Kit (Qiagen, Hilden, Germany). cDNA was synthesized using the iScript cDNA synthesis kit (Bio-Rad, Hercules, CA). qPCR was conducted using SsoAdvanced universal probes supermix (Bio-Rad, Hercules, CA) and target specific TaqMan real-time PCR assay primer probes (ThermoFisher Scientific, Waltham, MA). Viral burden was determined by quantitative real-time RT-PCR on lung RNA for viral matrix protein (M1) as described previously (67, 68). Forward Primer:5′-GGACTGCAGCGTAGACGCTT-3′, Reverse Primer:5′-CATCCTGTTGTATATGAGGCCCAT-3′,Probe:5′-/56- FAM/CTCAGTTAT/ZEN/TCTGCTGGTGCACTTGCCA/3IABkFQ/−3′.

### Protein Assays and Lincoplex

The Pierce^TM^ BCA protein assay kit (ThermoFisher Scientific, Waltham, MA) was used as directed to determine protein levels in BALF. Cytokine production was measured in lung homogenates via Bioplex using the Luminex™ Magpix™ multiplexing platform with the Bio-Plex Pro Mouse Cytokine 23-plex assay (Bio-Rad, Hercules, CA) as directed. Mouse lung homogenates were used to determine IgM protein using an IgM uncoated ELISA kit as directed (Invitrogen-ThermoFisher Scientific, Waltham, MA).

### Bulk-RNA Sequencing

RNA for sequencing was isolated from total mouse lungs as described above. Library preparation and sequencing was performed at UPMC Children’s Hospital of Pittsburgh. Samples were run on the Illumina NEXT-Seq 500 platform: 1x75bp single-end reads and 20 million reads per sample.

### Bioinformatics

Bulk-RNA seq reads were aligned using CLC Genomics Workbench 22 (Qiagen, Hilden, Germany). Once aligned, raw counts were extracted and exported to R workspace where the data was further processed with the DESEQ2 and clusterprofiler packages (69, 70). We used Cell-type Identification by Estimating Relative Subsets of RNA Transcripts, CibersortX, to estimate and define cell-types via RNA transcript information from our bulk-RNA memory versus acute super-infected sequencing data using a single-cell reference dataset (71, 72). In brief, a signature matrix was built from a publicly available single-cell dataset from PBS and 48-hour influenza infected mouse lung tissue that consisted of 6,528 cells (40) with 50% sampled without replacement for our 10 cell types of interest. Default parameters were used for the signature matrix building and CIBERSORTx analysis was conducted on our bulk-RNA sequencing dataset. Relative mode and number of permutations was set at 100 and cell type proportions, Pearson correlation coefficient, p value, and root mean squared error was calculated (RMSE). All sequencing data will be uploaded to Gene Expression Omnibus upon publication.

### Statistical Analysis

Data were analyzed using GraphPad Prism software (San Diego, CA). Experiments were repeated 3-6 times as indicated. All data are presented as mean <u>±</u> SEM, unless otherwise noted. Mann-Whitney test or one-way ANOVA followed by multiple comparisons were used for statistical significance with a p value of equal or less than 0.05. Mouse studies were repeated at least three times, in most cases.

## Results

### Preceding heterotypic influenza infection is protective against subsequent secondary bacterial infection

To determine the role that immune memory plays in influenza associated super-infections we primed wild-type (WT) C57BL/6 mice with influenza A/X31 H3N2 (X-31) on day zero followed by re-challenge with a heterotypic influenza strain, A/PR/8/34 H1N1 (PR8), and inoculation with methicillin-resistant *Staphylococcus aureus* USA300 (MRSA) six days after PR8 infection (Fig. 1a). Super-infected mice that had pre-existing heterotypic influenza immunity were better protected upon bacterial infection, with decreased weight loss and bacterial burden than their acute infection counterparts (Fig. 1b and c). When lung infiltrating cells were assessed, super-infected memory mice had lower levels of total infiltrating cells, and upon further assessment it was determined that monocytes made up more of the total cell distribution (Fig. 1d). Previous research on influenza associated bacterial super-infections have elucidated several potential mechanisms to understand the driving forces behind these types of infections with one of the prevailing ideas being that the anti-viral mechanisms in the lung environment dampen the anti-bacterial response (7, 9, 35, 36). We observed that memory super-infected mice at the time of MRSA challenge had non-detectable levels of PR8 gene expression in their lungs indicating that the anti-viral memory mechanisms controlled the infection, potentially ensuring a more effective anti-bacterial response to secondary bacterial infection (Fig. 1e). Finally, we observed a reduction in levels of Type I, II, and III interferons (Ifn), which are important signaling molecules during viral infection and have been shown to interfere with antibacterial response and survival outcome in secondary bacterial infections (17, 37–39) (Fig. 1f). These data suggest that mice with influenza memory have an advantage over their previously naïve acute infection counterparts upon challenge with MRSA due to a more controlled and targeted anti-viral response.

**Figure 1.**
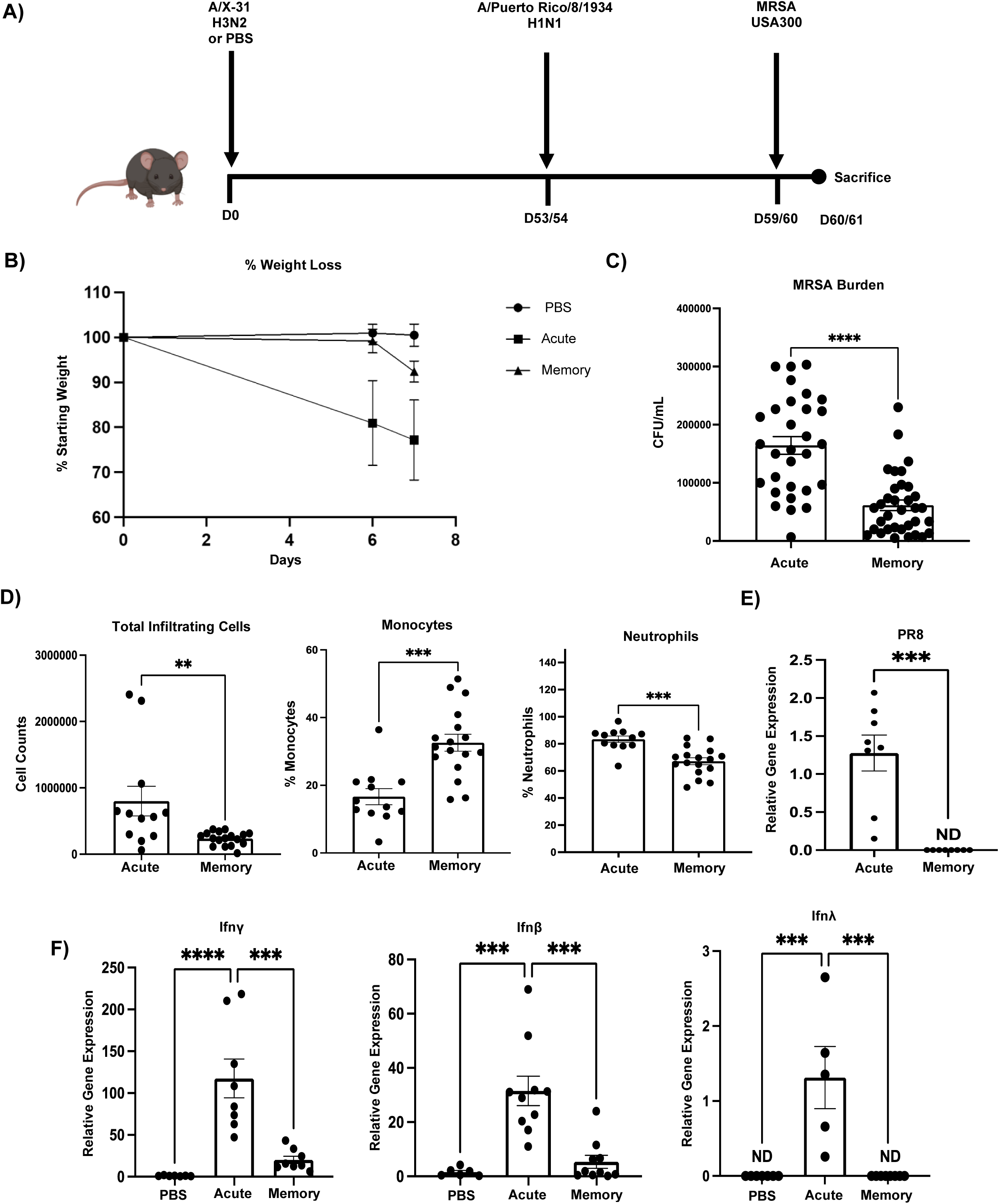
Influenza memory experienced mice are better protected against secondary bacterial infection with *Staphylococcus aureus.* A) Infection scheme for memory super-infected C57BL/6 mice. Mice were infected on day 0 with A/X-31 H3N2 (X-31) and at 53 to 54 days post infection (p.i.) mice were given A/Puerto Rico/8/1934 H1N1 (PR8). MRSA USA300 (MRSA) was given on day 59 or 60 p.i. and mice were harvested on day 60 or 61 p.i. B) Percent of weight loss was calculated starting from secondary infection with PR8 for each of the treatment groups and compared to mice that received PBS only, with day 6 being the date that MRSA was given on (PBS= 15, acute n=17, memory n=23). C) Number of MRSA colonies plated from infected mouse lung homogenates (acute n=30, memory n=34). D) Total infiltrating cell counts, percent of monocytes, as well as percent of neutrophils were determined by cytospins using bronchoalveolar lavage fluid collected from infected mouse lungs at time of harvest (acute n=12, memory n=16). E) Presence of viral protein was assessed in cDNA from total mouse lung via relative gene expression analysis of PR8 (Matrix protein) (acute n=8, memory n=8). F) Relative gene expression of interferon classes (Ifn-λ3, Ifn-γ, and Ifn-β) from total mouse lung cDNA. ND=non-detectable. p values: *<0.05, **<0.01 ***<0.001, ****<0.0001. Mouse figure was created with BioRender.com.

### Lung injury is present in acute and memory influenza challenged mice

Since viral burden was effectively controlled in heterotypic influenza challenged mice, we investigated if lung injury was altered in our model. Memory and acute super-infected mice were equally susceptible to lung damage following PR8 and MRSA challenge; however, the memory mice displayed areas of epithelial remodeling and metaplasia (Fig. 2a and b). Interestingly, when we looked at other measures to assess tissue integrity, we found that memory mice had attenuated levels of IgM and protein in their BALF when compared to their acute counterparts (Fig. 2c). The respiratory system has several mechanisms to defend itself against pathogens and to ensure a balance in the lung environment, because of these protective factors we also examined the expression of genes associated with lung function, protection, and remodeling. Memory experienced mice displayed elevated expression levels of important molecules involved in the mucociliary escalator: mucin 5b (muc5b) and mucin 5ac (muc5ac) (Fig. 2d). Further, secretoglobin family 1a member 1 (scgb1a1), a key club cell protein, and forkhead box j1 (foxj1), a key transcription factor in ciliated epithelial cells, were elevated in memory super-infection versus acute infection (Fig. 2e). Interestingly, we did see reductions in gene expression of epithelial remodeling markers in the memory mice most notably in collagen 1 a 1 (col1a1) and α – smooth muscle actin (acta2), but saw no change in tight junction protein (tjp1) (fig 2f). These data indicate that heterotypic memory to influenza drives changes in the lung environment that differ from their previously naïve acute counterparts and may hinder opportunistic bacterial infections.

**Figure 2.**
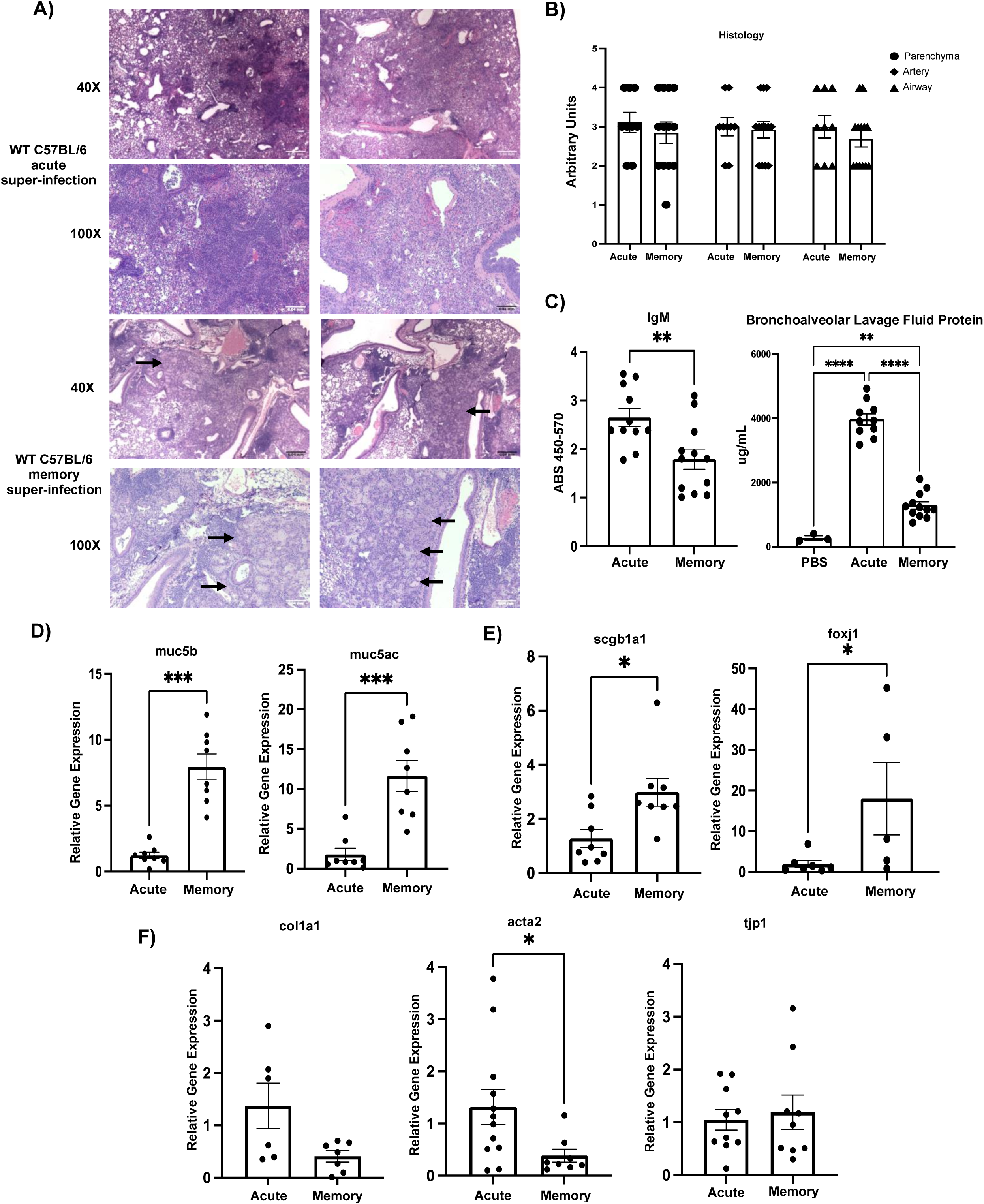
Lung damage is evident in acute and memory challenge. A) Representative histology sections of acute (n=2) and memory (n=2) super-infected WT C57Bl/6 mice from formalin inflated mouse lungs stained with H&E at 40X and 100X. Black arrows denote areas of metaplasia in memory mice. B) Blinded histology scores from paraffin embedded lung tissue sections. Lung injury scoring was carried out on three areas of the lung: parenchyma, artery, and airway (acute n=9, memory n=13).C) IgM was quantified from mouse lung homogenates using an IgM ELISA (acute n=11, memory n=12). BAL protein was quantified from BAL collected at day of harvest (PBS n=3, acute n= 10, memory n= 12). D-F) RNA was extracted from mouse lungs and made into cDNA which we then used to probe for various genetic targets (muc5b, muc5ac, scgb1a1: acute n=8, memory n=8; foxj1: PBS n=6,acute n=7, memory n=5; col1a1:acute n=6, memory n=7; acta2: acute n=12, memory n=8; tjp1: acute n=11, memory n=9). p values: *<0.05, **<0.01 ***<0.001, ****<0.0001.

### Lung transcriptomics indicate an altered inflammatory microenvironment and epithelial biology in influenza memory infected mice

To study the broad-spectrum changes in the lung environment of influenza memory super-infected mice and their previously naïve acute counterparts we compared gene expression changes in influenza memory versus acute infected mice via bulk-RNA sequencing analysis. We performed a deconvolution analysis on our day seven bulk-RNA sequencing dataset using a single-cell reference dataset of combined PBS and 48-hour influenza infected mice (40). Deconvolution analysis revealed changes in the proportions of genes associated with granulocytes, mononuclear phagocytic cells, epithelial cells, and α-sma between memory and acute infected groups (Fig. 3a). Influenza memory experienced mice had a transcriptomic signature driven more by epithelial cells and less by granulocytes compared with acute infection controls. Sequencing analysis showed clustering of samples within groups (Fig. 3b). When up or down-regulated genes were assessed in memory versus acute infection we found that memory mice displayed a reduction in genes typically associated with inflammation and control of inflammatory mechanisms such as interferon gamma (Ifn-γ), granzyme B (Gzmb), and interleukin (IL)-10 (Fig. 3c). Next, we ran gene ontology analysis on our sample groups and found that the pathways activated in memory are associated with the epithelium and mucociliary mechanisms, whereas those that are suppressed deal with response to virus, and immune effector processes as well as the inflammatory response (Fig. 3d). Gene ontology biological pathway analyses revealed that the top enriched pathways dealt with leukocyte migration, response to interferon-beta, response to chemokine, as well as chemotaxis (Fig. 3e). Together these data demonstrate that the controlled lung environment of influenza memory experienced mice is characterized by changes towards a state of lower inflammation and a balance in inflammatory mechanisms potentially accounting for a better immune response to a secondary bacterial infection.

**Figure 3.**
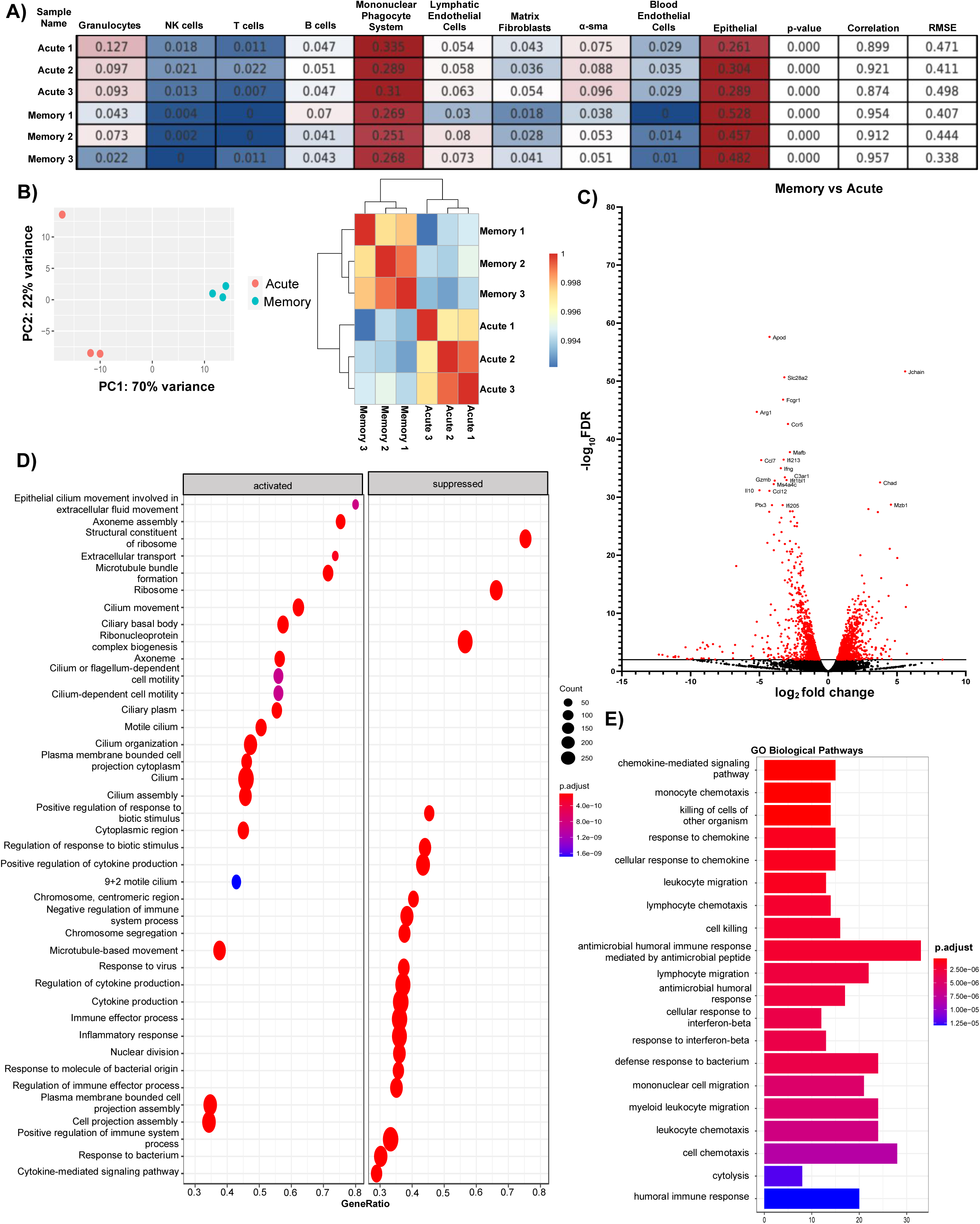
The immune signature of memory experienced mice shows a shift in inflammatory pathways and highlights a trend towards a less inflamed lung environment. Illumina reads were aligned using CLC Genomics Workbench 22 and raw counts were extracted for bulk RNA seq analysis using DESEQ2 and clusterprofiler R software packages (acute n=3, memory n=3). A) CibersortX was used to perform deconvolution on bulk-RNA sequencing data using a single reference dataset that contained 48-hour influenza and PBS-treated mouse lung data. Numbers in boxes reflect proportion of cells in each group. B) PCA plot and clustering heatmap from Bulk RNA-seq analysis showing how the acute and memory infected groups cluster. C) Volcano plot of differentially expressed genes in memory versus acute infected mice. Red dots denote genes with a -log_10_FDR >2. Genes highlighted are the top 20 genes with - log_10_FDR >2. D) Gene Ontology analysis of activated and suppressed pathways in memory vs acute infected mice with p values>0.05. E) Gene enrichment analysis on biological pathways in memory versus acute infected mice on genes with p values>0.05.

### Heterotypic influenza memory alters the lung T cell compartment during bacterial super-infection

To further characterize the changes in the mouse lung inflammatory environment driven by the presence of pre-existing influenza memory, we examined pulmonary immune cell subsets. T cell populations play a vital role in clearance of influenza infections, therefore, we looked at how the T cell compartment changes in influenza memory versus acute super-infected mice. Using clustering (flowSOM) and dimensionality reduction (UMAP) methods, we tracked marker expression changes in T cell populations in mouse lungs. From our gating strategy we found that (Supplementary Fig 1), in an acute super-infection there are a higher proportion of CD4^+^ FoxP3^+^ regulatory T cells, this significant change was seen in both percentage of parent gate as well as absolute cell counts (Fig. 4a-c). The shift towards a lower proportion of regulatory T cells in the memory lung environment could indicate an earlier controlled inflammatory response towards viral infection allowing for an unimpeded switch towards immune mechanisms generated in response to bacterial infections. Further, we observed an increase in the proportion of CD8^+^ NP^+^ tetramer-positive T cells, an immunodominant epitope, in memory experienced mice when compared to the acute infected mice. This was again confirmed by significant differences in cell proportions and absolute cell counts consistent with increased memory expansion (Fig. 4a, b, d). We next looked at the expression of cytokines associated with T cell migration, proliferation, and regulation. We found that protein levels of pro-inflammatory cytokines IL-12p40 and IL-12p70 were significantly elevated in influenza memory experienced super-infected mouse lungs (Fig. 5a). Other cytokines that significantly changed in the lungs of memory or acute super-infected mice were cxcl10, IL-6, and IL-10 all of which are significantly decreased in memory experienced mice via either protein expression or relative gene expression (Fig. 5b-d). Lastly, we saw a trend toward higher levels of IL-17a and IL-22 via gene expression, although this was not significant, however, we did see a significant increase in expression of the Type 17 immune promoting cytokines IL-23 and IL-1β (Fig. 5e). These data demonstrate that the T cell compartment in memory experienced mice is characterized by an influx of tetramer positive CD8^+^ T cells that are better able to control influenza infection, thus creating an environment that is potentially oriented towards improved bacterial immunity.

**Figure 4.**
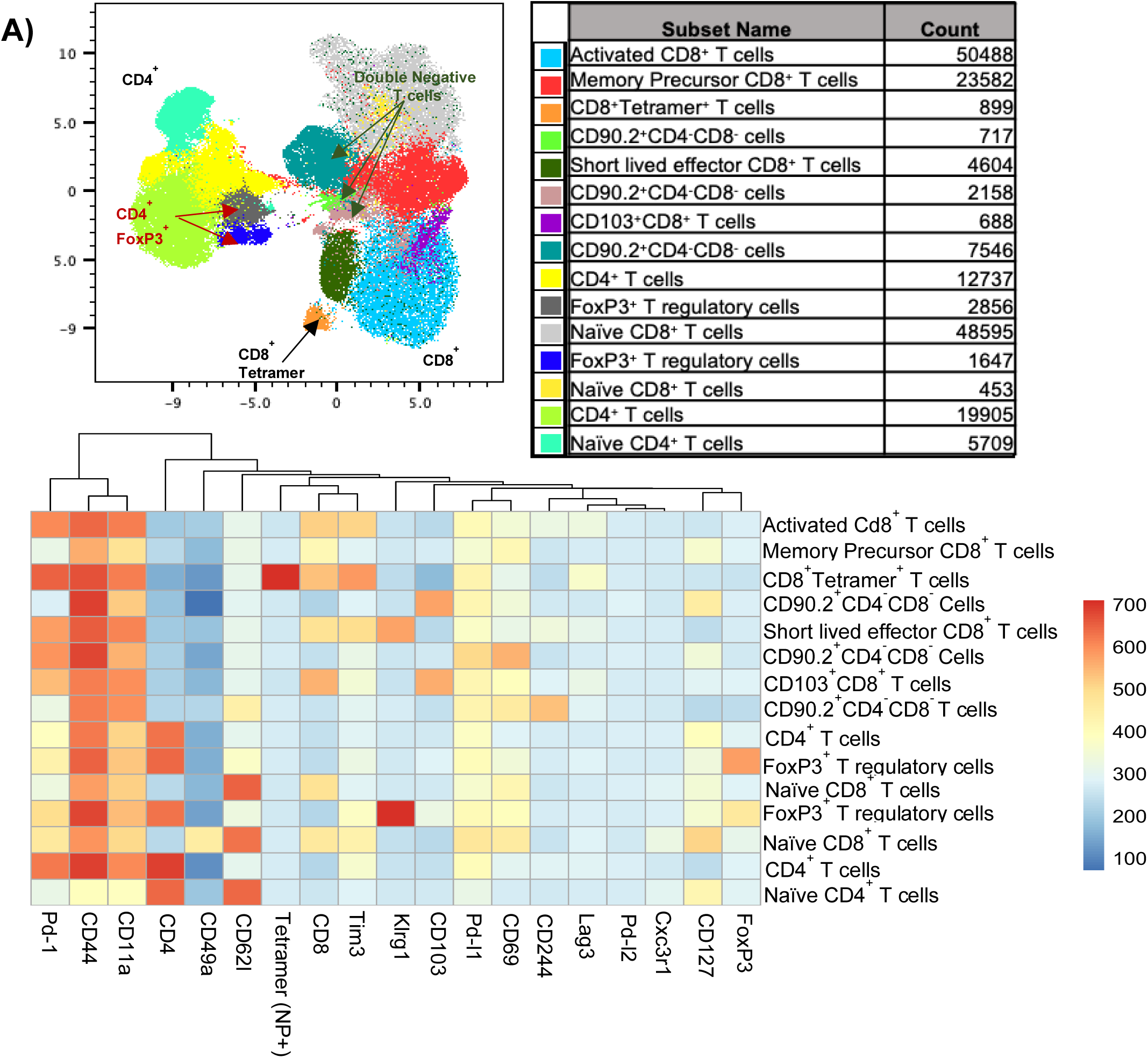
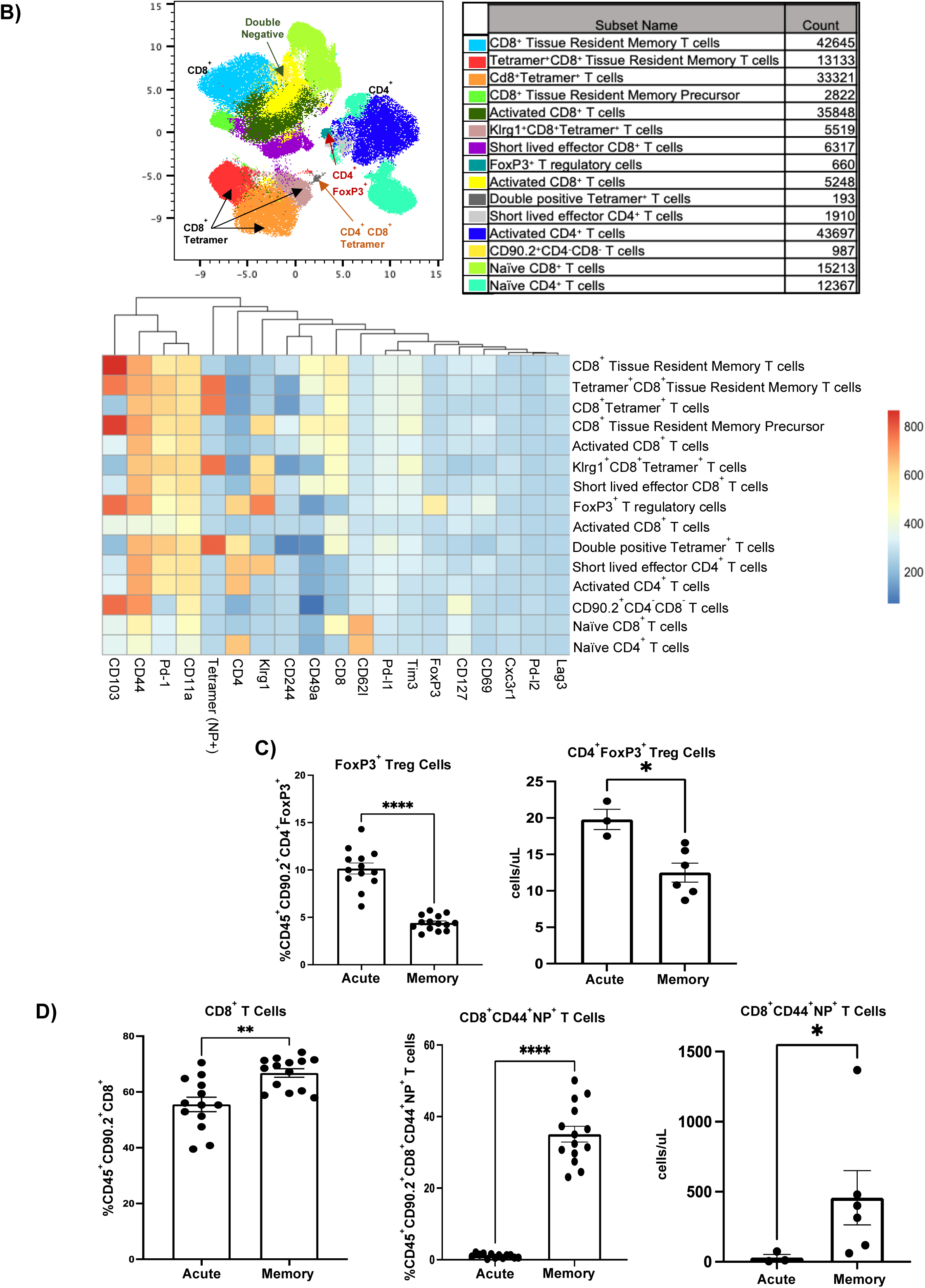
Memory experienced mice have a distinct T cell landscape characterized by an increase in NP+ tetramer specific CD8^+^ T cells. A) Flow cytometry analysis on previously naïve acute super-infected mouse lungs. Samples were initially gated on CD45^+^ CD90.2^+^ Live cells. Once gated samples were concatenated (n=4) and populations were visualized using FlowSOM and UMAP plugins in FlowJo. FlowSOM populations were further analyzed by conventional gating techniques to determine breakdown of T cell types. B) Flow cytometry analysis on memory super-infected mouse lungs (n=5). Gating and visualization of T cell types were determined as noted above for A. C) Percentage and absolute cell count of CD45^+^ CD90.2^+^ CD4^+^ FoxP3^+^ Treg cells determined from flow cytometry in mouse lungs (percentage FoxP3^+^ acute n=13, memory n=14, absolute number FoxP3^+^ acute n=3, memory n=6). D) Percentage of CD45^+^ CD90.2^+^ CD8^+^ T cells and NP^+^ T cells with absolute cell counts determined by flow cytometry in mouse lungs (percentage acute n=13, memory n=14, absolute cell counts acute n=3, memory n=6). p values: *<0.05, **<0.01 ***<0.001, ****<0.0001.

**Figure 5.**
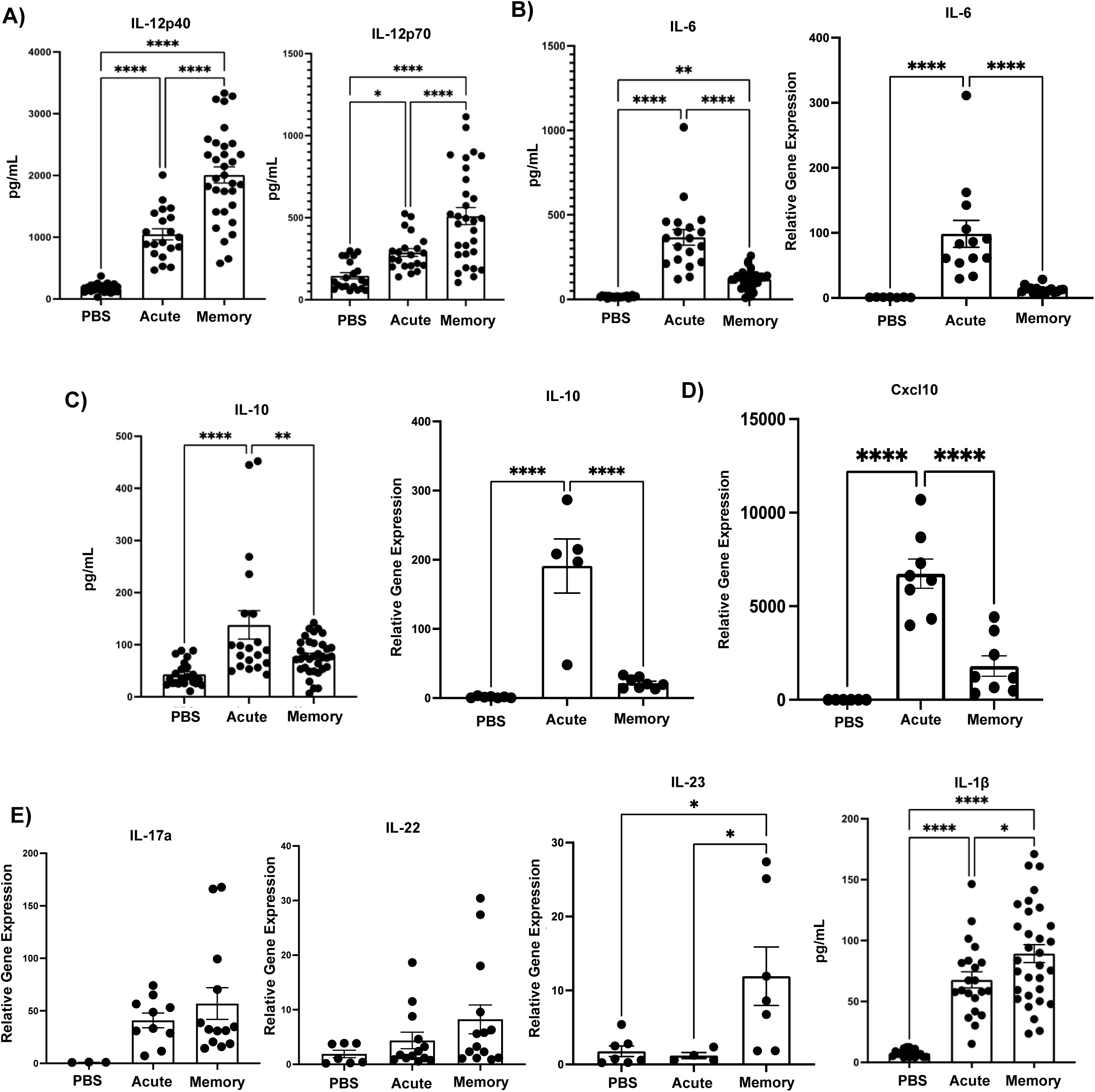
Memory experienced mice have reduced levels of cytokines that act as indicators of severe influenza infection, but display elevated levels of cytokines with roles in bacterial clearance. A-E) Relative gene expression and protein levels in memory versus acute super-infected mouse lungs. (IL-12p40 PBS n=23,acute n=20, memory n=32; IL-12p70 PBS n=21, acute n=21, memory n=30; IL-6 PBS n=21, acute n=19, memory n=29; relative gene expression IL-6 PBS n=7, acute n=13, memory n=16; IL-10 PBS n=22, acute n=20, memory n=34; relative gene expression cxcl10 PBS n=6, acute n=8, memory n=8; relative gene expression Il-17a PBS n=3, acute n=10, memory n=13; relative gene expression IL-22 PBS n=7, acute n=13, memory n=14; relative gene expression IL-23 PBS n=7, acute n=4, memory n=7; IL-1β PBS n=18, acute n=21, memory n=31). p values: *<0.05, **<0.01 ***<0.001, ****<0.0001.

### Heterotypic influenza memory alters the innate immune cell compartment during bacterial super-infection

Finally, we explored the impact of heterotypic influenza memory on innate immune cell populations, which are imperative in defense against bacterial infections. We used flowSOM and UMAPs to visualize changes in the myeloid compartment when mice have an established memory compartment. We found that neutrophils make up the largest proportion of myeloid cells in both treatment groups (Fig. 6a and b). Using our myeloid gating strategy (Supplementary Fig 2), we also found that the total number of natural killer (NK) cells, monocytes, macrophages, and eosinophils were significantly reduced in the influenza memory mice as compared to their acute counterparts (Fig. 6c). Interestingly, no change was seen in the total number of neutrophils between both groups (Fig. 6c). Next, we assessed changes in common cytokines and markers associated with myeloid cells and their function via relative gene expression as well as protein expression. We first looked at two mediators of the inflammatory response, the alarmins IL-1α and IL-33. Both of these cytokines were found to be significantly elevated in the lungs of memory mice via protein expression or relative gene expression (Fig. 6d). When we looked at markers that were significantly downregulated in the influenza memory mice we observed a decrease in protein expression of monocyte chemoattractant protein-1 (mcp-1), macrophage inflammatory protein-1 beta (mip-1β), macrophage inflammatory protein-1 alpha (mip-1α), and eotaxin (Fig. 6e), which suggests a less inflamed environment. Lastly, we found that arginase-1 (arg1), cathepsin g (ctsg), nitric oxide synthase 2 (nos2), and amphiregulin (areg) expression were also downregulated in memory mice (Fig. 6f). The promotion of an inflammatory balance in the lung environment through the stages of infection from influenza to secondary bacterial infection in the heterotypic memory mice may create an environment that is suitable for bacterial clearance.

**Figure 6.**
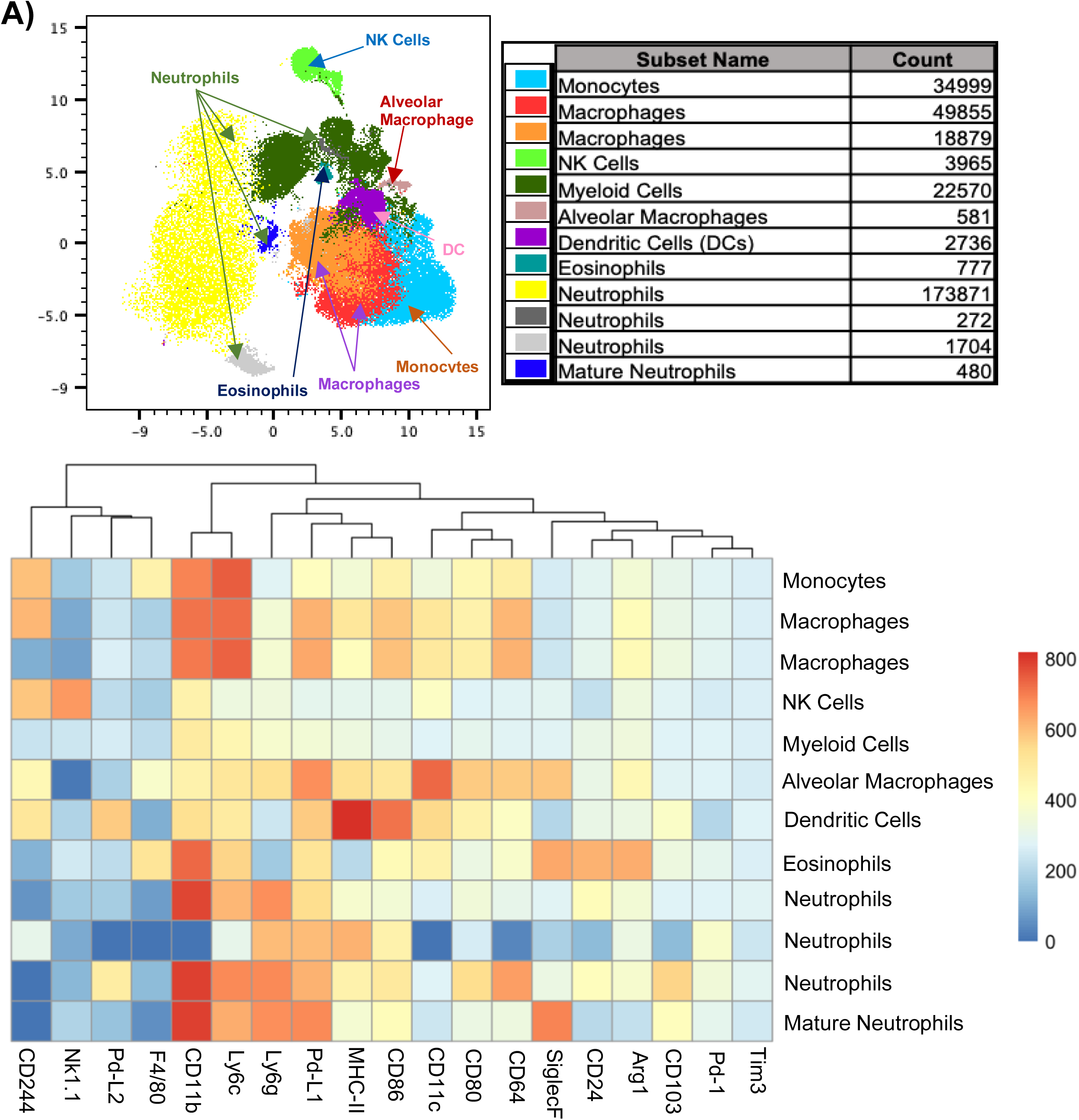
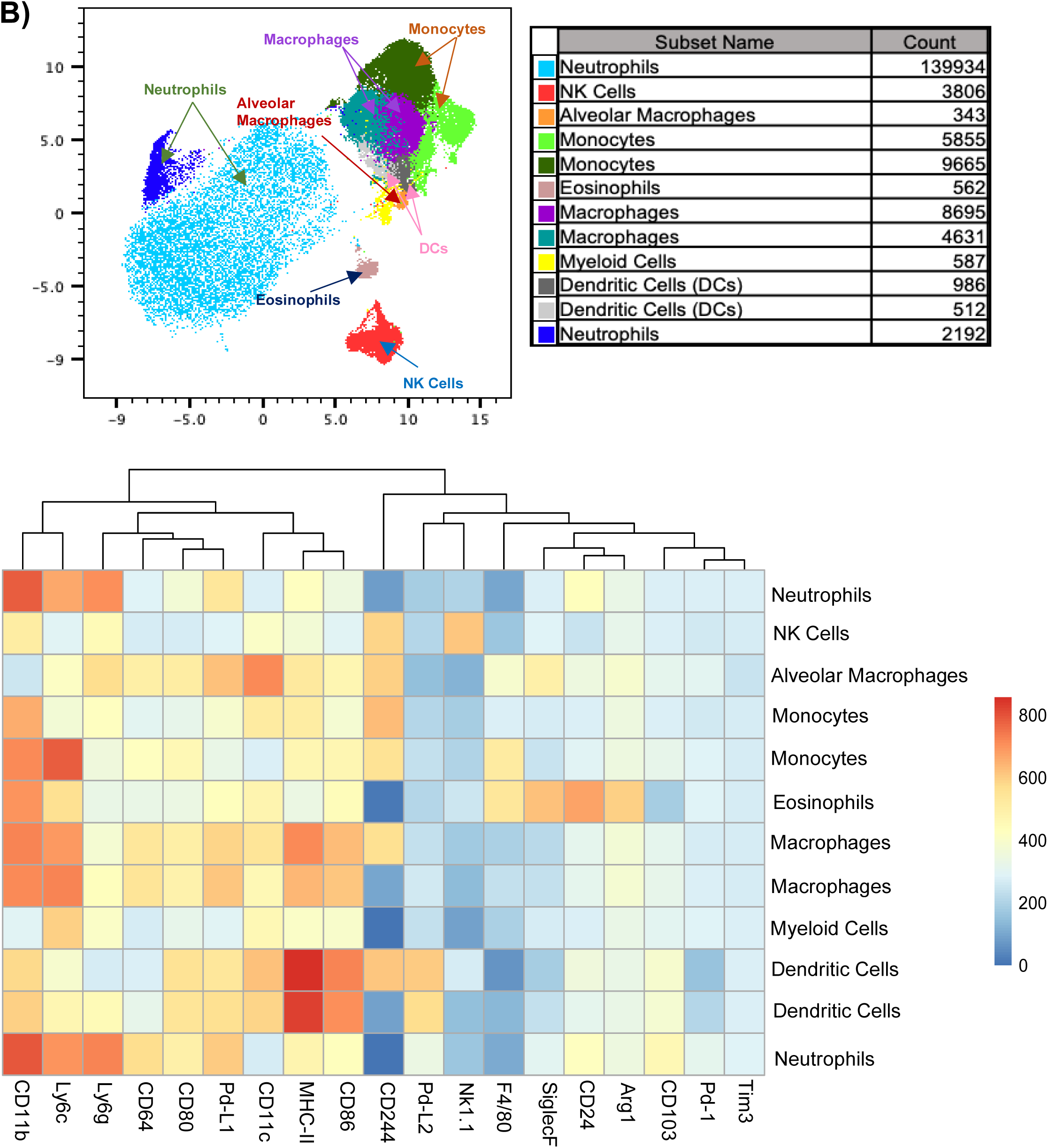
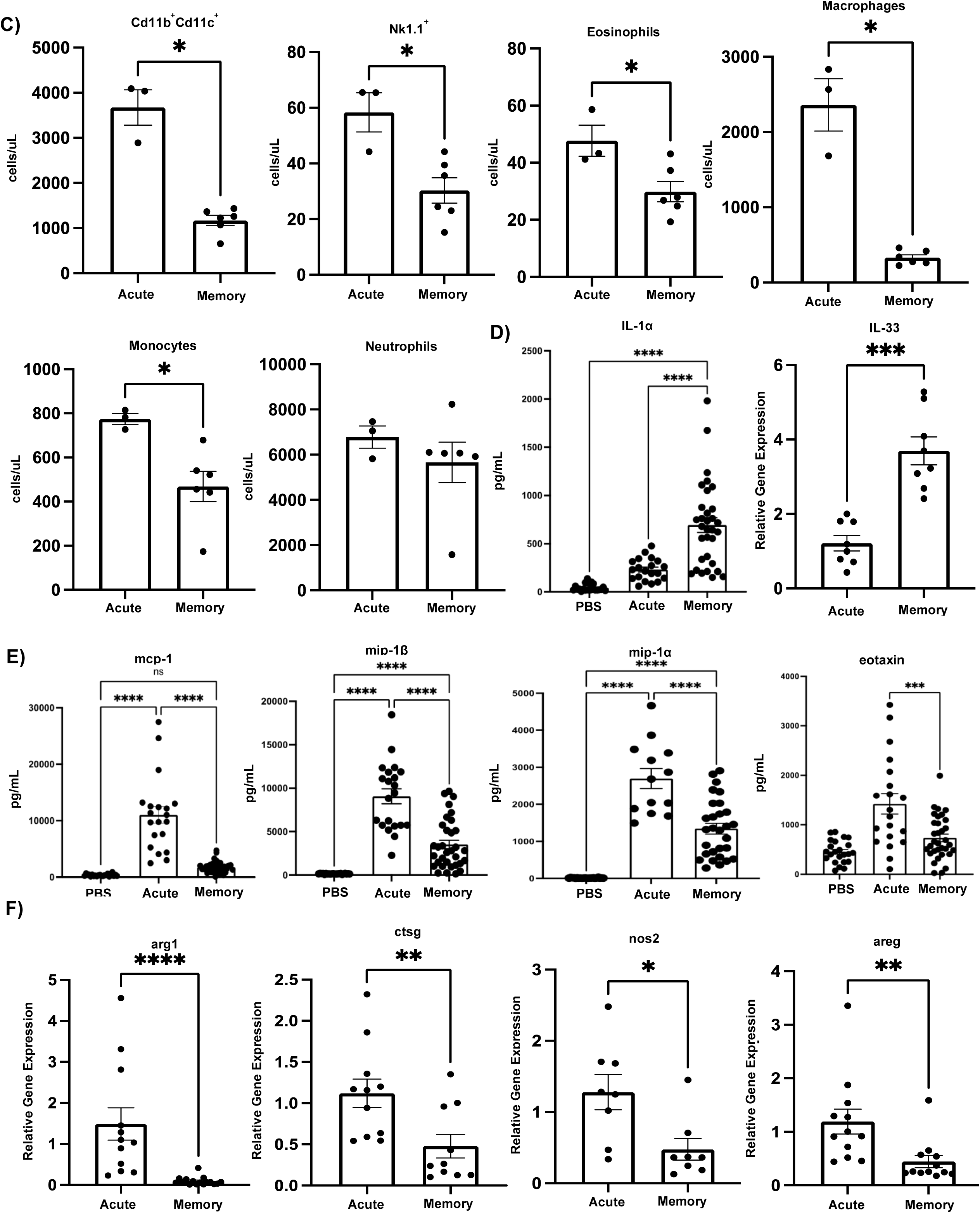
The myeloid cell landscape in memory experienced mice is characterized by a reduction in the number of cell types. A) Flow cytometry analysis on acute super-infected mouse lungs. Samples were initially gated on CD45^+^ TCRβ^-^B220^-^Live cells. Once gated samples were concatenated (n=5) then downsampled and populations were visualized using FlowSOM and UMAP plugins in FlowJo. FlowSOM populations were further analyzed by conventional gating techniques to determine breakdown of myeloid cell types. B) Flow cytometry analysis on memory super-infected mouse lungs (n=4). Gating and visualization were determined as noted above for A. C) Absolute cell counts of myeloid cell subsets from flow cytometry analysis (acute n=3, memory n=6). D) Protein expression of IL-1α (PBS n=22, acute n=20, memory n=32) and relative gene expression of IL-33 (acute n=8, memory n=8) in mouse lungs. E) Protein expression of cytokines associated with the myeloid compartment (mcp-1 PBS n=22, acute n=20, memory n=34; mip-1β PBS n=22, acute n=21, memory n=33; mip-1α PBS n=22, acute n=13, memory n=29; eotaxin PBS n=23, acute n=20, memory n=33). F) Relative gene expression of molecules associated with myeloid cell function (arg1 acute n=12, memory n=16; ctsg acute n=11, memory n=10; nos2 acute n=8, memory n=8; areg acute n=12, memory n=12). p values: *<0.05, **<0.01 ***<0.001, ****<0.0001.

## Discussion

The importance of understanding how bacterial and viral infections synergize together is widely applicable across many respiratory viral diseases including the recent Covid-19 pandemic (41). Influenza associated secondary bacterial infections are multi-faceted infections that employ multiple immune cell players of both the adaptive and innate immune systems to cover the broad spectrum of the host response from viral infection to bacterial infection. It is widely established that during an acute secondary bacterial infection the anti-viral response hinders the anti-bacterial mechanisms that are required to protect against opportunistic pathogens such as MRSA (7, 9, 12, 18–20, 36, 39, 42–46). Secondary bacterial infection also results in heightened immunopathology driven by an overactive immune landscape ultimately creating an ideal environment for opportunistic pathogens (13). The results presented herein expand on the prevailing views in the field generated in previously naive animals, and suggest that memory to influenza infections creates a balanced lung environment at the time of secondary bacterial infection susceptibility resulting in a more effective anti-bacterial response.

The epithelium represents the first line of defense against respiratory pathogens and assault from environmental stressors. Influenza directly targets the epithelial barrier leading to extensive damage and impairment of the mucocilliary escalator resulting in an environment that heavily favors bacterial adherence and colonization (42, 47). In the present study we found that influenza memory experienced mice have an altered lung epithelial environment as compared to their previously naïve acute infection counterparts. Interestingly, our data shows that the influenza memory lung still undergoes significant damage due to influenza infection. However, we observed preservation of epithelial gene expression of structural and functional markers in influenza memory mouse lungs. In our data, we saw an increase in the expression of Scgb1a1 (also known as club cell secretory protein), which has been shown to promote alveolar macrophage survival and response to inflammation (48). Deconvolution analysis also suggested that memory mice expressed a higher proportion of the lung gene signature associated with the epithelium. Further, we observed upregulation of markers associated with cilium and ciliary movement, as well as mucus production. Taken together these data suggest that influenza memory doesn’t prevent lung injury, rather the epithelium is re-programmed in a manner that favors the enhancement of bacterial clearance. Barrier integrity is also targeted by aberrant immune mechanisms including enhanced cytokine production in response to influenza that in cases of severe viral infections can cause severe edema (49). Lung leak was also limited in the context of heterotypic influenza memory. Aberrant immune activation, such as cytokine storm, is key to influenza related lung injury. Our data suggests that there is a reduction in production of cytokines that are detrimental and associated with immunopathology, in particular mcp-1, mip-1α, cxcl10, and IL-6 (49–52).

A critical immune response to viral infection is the production of interferons, a highly inducible first line defense against influenza infection. The interferon group is made up primarily of three classes: type I (Ifn-α and Ifn-β), type II (Ifn-γ), and type III (Ifn-λ). Studies examining the role of interferons during bacterial super-infection have shown that they enhance susceptibility. The mechanisms highlighted in these studies are due to decreased Type 17 immune activation and destabilization in function and recruitment of phagocytic cells (10, 11, 14, 17, 39, 53, 54). Our data suggests that at the time of super-infection, influenza memory experienced mice have reduced levels in expression of interferons, which based on prior studies would indicate that anti-bacterial mechanisms are preserved. One of the major roles that interferons play in the immune system is to induce production of other chemokines such as cxcl10, an important molecule associated with the trafficking of immune cell types including activated T cells (52). Although, this immune cell trafficking to the site of infection is imperative for resolution of infection, too many pro-inflammatory cells in the lung environment can lead to detrimental outcomes. In this study, we observed a decrease in the relative gene expression of cxcl10 that could potentially indicate decreased inflammation in the lung environment of memory experienced mice, and adds further support to the reduction seen in infiltrating cells at time of super-infection.

An effective anti-bacterial response is marked by the presence and activation of phagocytic cells. Support for a more effective anti-bacterial response to MRSA infection in our heterotypic memory experienced super-infected mice is demonstrated in the increase in the IL-1 family of cytokines, including IL-1β, IL-1α, and IL-33. It has been shown that restoration of IL-33 during an influenza associated bacterial super-infection is beneficial for bacterial clearance of both *S. aureus* as well as MRSA due to the role it plays in neutrophil recruitment to the infected lung (55). IL-1β production has been shown to be impaired by preceding influenza infection during secondary bacterial super-infections. This impairment results in a detrimental outcome for the host because of its ability to influence and activate the Type 17 pathway, an important pathway during antibacterial response against numerous extracellular pathogens including *S. aureus* (56). The IL-12 family has also been implicated in its role in protection against bacterial infections. In our model, we found that three members of the IL-12 family: IL-12p40, IL-12p70 and IL-23 were increased. IL-12p70 and IL-23 have subsequently been shown to play a role in bacterial super-infections in human cells due to their alteration by influenza resulting in impairment of the Ifn-γ and Type 17 pathways, respectively (57). Another study suggested that IL-12p40 is crucial to early protection against bacterial infections due to its role in neutrophil recruitment (58). However, this study also determined that Ifn-γ was important in their model and we saw neither an increase in the relative gene expression of Ifn-γ or neutrophil recruitment in our memory mice at the time of super-infection. Although, we did not see a significant increase in Type 17 cytokines, IL-17a and IL-22 in our influenza memory versus acute infected mice, we did see increased IL-1β, IL-23 and decreased Ifn-γ, suggesting promotion of Type 17 immunity. Together these data indicate that the heterotypic memory mice display increases in pathways related to bacterial defense and further studies on these pathways should be done to elucidate direct mechanisms.

The lung is a dynamic organ that requires a state of homeostasis characterized by a balance in pro-inflammatory and anti-inflammatory factors (59). Influenza infections result in a highly inflamed lung environment that is characterized by an influx in immune cells and their mediators. To return to homeostasis, the lung initiates anti-inflammatory mechanisms. One of the key primary anti-inflammatory mediators of the immune system is Foxp3^+^ regulatory T cells. These cells expand in response to an overly inflamed mucosal environment to dampen the immune response and have been shown to acquire memory recall for targeted pathogen responses during re-challenge events (60, 61) which are imperative to bring the inflamed environment back to homeostasis. In our study we saw a reduction in the proportion and counts of FoxP3^+^ regulatory T cells in heterotypic memory experienced mice, as well as a reduction in the production of the anti-inflammatory cytokine IL-10. IL-10 has further been implicated in impairment of immune defense against influenza associated secondary bacterial infections with *Streptococcus pneumoniae* and *Staphylococcus aureus* (62–64). These data suggest that the lung environment of the influenza memory mice is targeted against the virus early on in re-challenge, which is reflected in the lower number of regulatory T cells present at time of bacterial infection. Our data further supports this notion with the increase seen in tetramer specific T cells in memory mice and reduction in innate immune cells that are elevated in acute infections and are associated with aberrant pro-inflammatory responses.

In order to promote an environment that is beneficial for bacterial clearance and limits immunopathology caused by aberrant immune activation, a balancing act must be struck between anti-viral and anti-bacterial host defense. As shown herein, this balance can be accomplished in memory experienced mice by a quick and controlled response against the influenza infection ultimately limiting the deleterious impact of anti-viral mechanisms on anti-bacterial immunity. This finding is highly relevant to the human condition where nearly all humans have some level of pre-existing immunity against influenza and is suggestive of an important role for pre-existing heterotypic influenza memory in limiting super-infection susceptibility. Further research on the role that heterotypic memory plays to enable susceptibility or protection against secondary bacterial infections is necessary and could provide further avenues of therapeutic intervention.

## Acknowledgements

We would like to thank Drs. Yael Steurman and Irit Gat-Viks (Tel Aviv University, Israel) who provided us with an annotated single cell reference dataset from their published data for the deconvolution analysis. We would also like to thank the Unified Flow Core at the University of Pittsburgh which houses and maintains the Cytek Aurora instruments (Cytek Biosciences, Fremont, CA). Lastly, this project used the University of Pittsburgh Health Sciences Sequencing Core at UPMC Children’s Hospital of Pittsburgh, Illumina NEXT-Seq platform.

