## Supplemental Material for "Heterotypic Influenza Infections Mitigate Susceptibility to Secondary Bacterial Infection"

**Supplementary Figure 1**

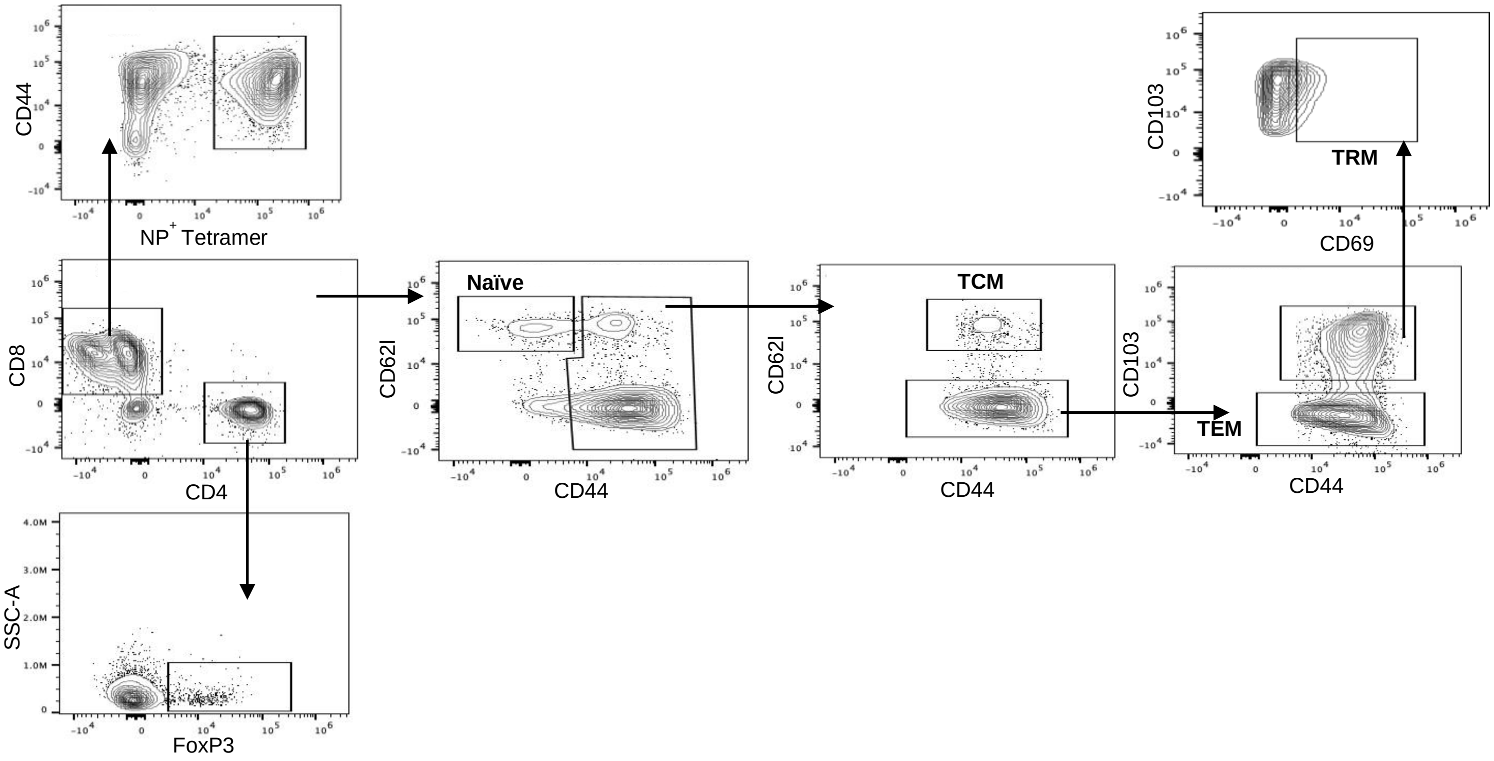


**Supplementary Figure 1. Gating strategy for T cell compartment analysis.** T cells were initially gated on CD90.2$^+$CD45$^+$Live cells and further gated to parse out changes in CD4$^+$ and CD8$^+$ T cells.

### Supplementary Figure 2

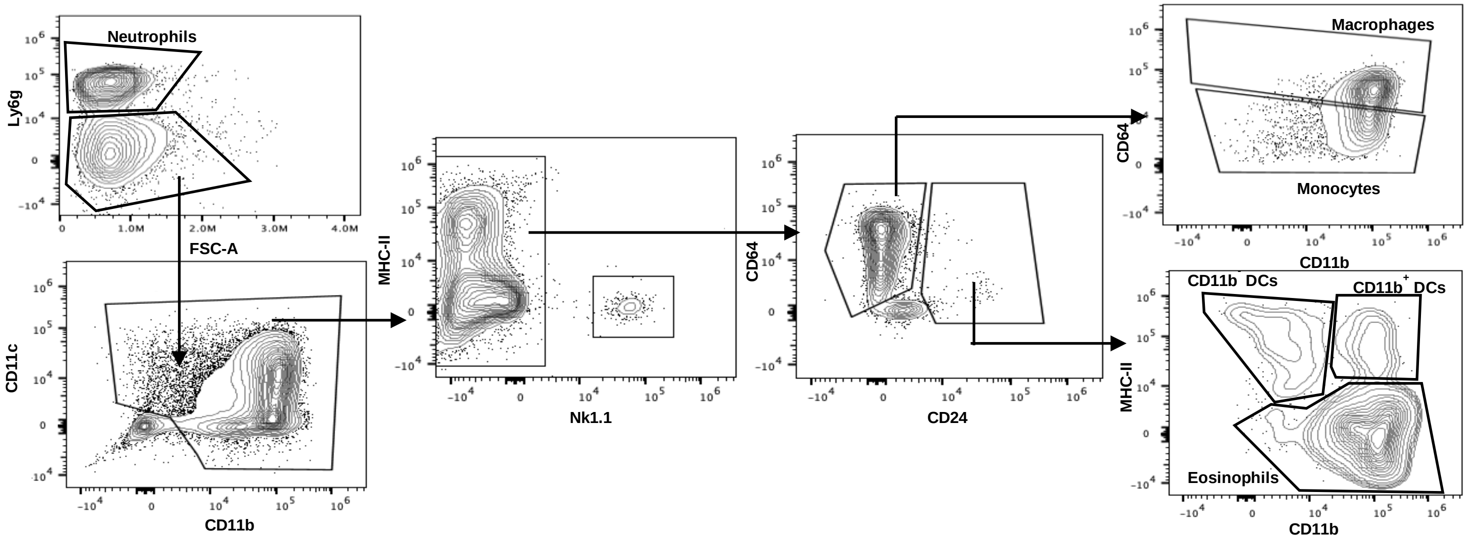


**Supplementary Figure 2. Gating strategy for myeloid cell subset analysis.** Myeloid cells were initially gated on TCRB$^{-}$B220$^{-}$CD45$^{+}$Live cells and further gated to determine changes in myeloid cell subsets.
